# Effects of land-use and landscape drivers in the species richness and distribution of carnivores in Faragosa-Fura Landscape of Southern Rift Valley, Ethiopia

**DOI:** 10.1101/2021.08.12.456157

**Authors:** Berhanu Gebo, Serekebirhan Takele, Simon Shibru

## Abstract

Understanding the species richness and distribution of carnivores across anthropogenic land-use types in an area is an essential first step for biodiversity conservation and human-carnivore coexistence. However, quantitative data on carnivore species coexisting with humans in different land-use types remain largely missing. Thus, this paper investigated the effect of anthropogenic land-use and landscape drivers on carnivore species richness and distribution in the Faragosa-Fura Landscape, Gamo Zone, southern Ethiopia. To collect data, we employed the line transect method using three complementary field surveys techniques: sign survey, camera-trapping, and opportunistic sighting survey during wet and dry seasons in 2020 and 2021. We stratified the study landscape into five land-use types-forest, wetland, grassland, agricultural land, and settlement. The result proved the occurrence of 12 carnivore species belonging to six families, including vulnerable Felidae species - *Panthera pardus*. Family Felidae and Herpestidae were composed of a greater number of species, while Hyaenidae and Mustelidae were each represented by single species. Out of identified species, only two species (*Panthera pardus* and *Crocuta crocuta*) were large-sized, while the rest were medium and small-sized carnivores. Overall, the mean richness of the study area was 5.73±0.284(SE). The species richness was highest in the wetland (n = 12, mean = 7.67±0.494(SE)) and lowest in the settlement (n = 5, mean = 4.25±0.479(SE)). The regression analysis showed that most of the carnivores displayed a strong negative relationship with agriculture, roads, and settlement while displayed a strong positive relationship with wetland and forest. In general, out of 32 species recorded in Ethiopia, this study quantified 12 carnivore species that signify the area is an important area for wildlife conservation in Ethiopia. Further, the study concluded that the wetland is the most important habitat, particularly for larger-sized and habitat specialists while anthropogenic land-uses types adversely affecting species richness. Thus, a generic paradigm to reconcile land management and biodiversity conservation is highly important.

## Introduction

Order Carnivora is composed of 290 species belonging to 16 families. Of these, 72 and 32 different species have been found in Africa [1], and Ethiopia [2, 3], respectively. Felidae, Viverridae, Herpestidae, Hyaenidae, Canidae, and Mustelidae are the six families of mammalian carnivores identified in Ethiopia where the 31 carnivore species belongs [2]. Ethiopia hosts two of the world’s 34 biodiversity hotspots [3]. However, carnivore species richness and distribution are being affected largely due to anthropogenic land use such as agriculture, settlement, and road infrastructure development [4]. For instance, the endemic Ethiopian wolf is among the most endangered carnivore species in the world [5]. In addition, African lions and leopards are under threat of local extinction at different localities [3]. Therefore, a survey on carnivore species richness and distribution in land-use types is vital for wildlife conservation.

The two most common strategies to conserve carnivores are establishing protected areas to separate carnivores from human-dominated areas [6] and promoting human-carnivore coexistence (HCCo): a sustainable state in which humans and wildlife co-adapt to living in shared landscapes [7, 8]. However, protected areas and native forests are diminishing largely due to anthropogenic loss of habitat, forcing carnivores to local extinction and/or to share habitats with humans [5, 9, 10]. This habitat sharing, coexistence, is resulting in human-carnivore conflicts that affect both local community livelihood and biodiversity conservation [8, 11]. About 95% of ranges of carnivore species occur outside protected areas in a human-dominated landscape [12]. Thus, understanding the carnivore species living with humans is important for both carnivore conservation and the livelihood of the local community [13, 14].

Therefore, successful conservation of carnivores must focus upon both in protected areas and a human-dominated landscape [6, 15]. Furthermore, the human-dominated landscape can also be very important for medium to small-sized carnivore conservation [12, 13, 16]. Therefore, habitats within a human-dominated landscape are increasingly important for carnivore conservation since protected areas are diminishing at an alarming rate.

Recent studies revealed that anthropogenic and landscape factors can potentially affect the carnivore species richness and distribution in different habitats. The influential factors are land-use types such as forest, grassland, wetland, agricultural land, and human settlement [11] because of differences in resources. While agriculture is the backbone of socio-economic development in developing countries like Ethiopia, it often comes at the expense of biodiversity and ecosystem services [17, 18]. Further, altitude and distance from the road [15, 19] are landscape factors affecting carnivore species richness and distribution.

Understanding the species richness (number of species) and species composition in an area is an essential first step for biodiversity conservation. These two biodiversity attributes have been mostly used in assessing the wildlife conservation potential of areas and devising the strategies of conservation efforts. The higher number of species and groups indicates a healthier community because a greater variety of species leading to greater system stability [3, 12].

At the community level, carnivores have been well studied in different parts of the world [6]. Nevertheless, in Africa carnivore surveys focused on single species rather than at the community level [15, 16]. The same is true in Ethiopia that no quantitative studies have been carried out on carnivores at a community level, but there exist only at the species-specific level [18, 20]. For instance, species-specific studies conducted for lion and hyena [21, 22], leopard [23], African civet [24, 25], mongooses [26], and Ethiopian wolf [27].

In Ethiopia, most protected areas are too small to sustain wide-ranging carnivore species [3]. As a result, carnivores are shifting to a human-dominated landscape and coexisting with humans that call for biodiversity conservation efforts in shared landscapes. However, the study of coexisting carnivore species with humans at a community level in the human-dominated landscape was missed largely.

We carried out this study in the Mirab Abaya district in a Faragosa-Fura Landscape (hereafter FFL) Gamo Zone, Southern Ethiopia. The study landscape is heterogeneous and largely a forest area that harbors distinctive carnivore species. The study area is associated with Lake Abaya, the largest lake in the Ethiopian Rift Valley system, which is the main water source for carnivore species and the lake-created wetland habitat. Despite this, the landscape has been under human-induced pressures. These pressures are adversely affecting the wildlife of the landscape. Thus, understanding the species richness and distribution of carnivore species in the area is important to urgent management actions. Moreover, there are no earlier ecological studies carried out in the study area vis-à-vis carnivores’. We examined the effects of land use and landscape drivers on the species richness and distribution of carnivores in this landscape. The central premise of this study is that the species richness and distribution of carnivores would be adversely affected by land-use types (agriculture, roads, settlement), the factors that fragment and degrade habitat. On the other hand, we expected natural forest cover (wetland, forest, grassland) to be critical in maintaining species richness and perhaps distribution of carnivores in the study landscape. Although the effect of landscape drivers (altitude, water source) might not affect the carnivore community, however, could drastically compromise the species’ ability to cope with the change. Our study quantifies the effects of land use and landscape drivers in the carnivore community of the hitherto understudied in the Faragosa-Fura Landscape of Southern Rift Valley, Ethiopia. Based on the finding, we recommend a generic paradigm to reconcile land management (land sharing, land sparing) and biodiversity conservation.

## Material and Methods

### Ethics Statement

All permissions to carry out field research were obtained from the Office of the environmental protection, Mirab Abaya district of southern Ethiopia. The Ethiopian Wildlife Conservation Authority’s (EWCA’s) Policy for Management of Wildlife Resources guidelines have been followed. Wherever observational investigations were made during walk transects, no animals were harmed.

### Study site

The study was conducted in the FFL that is found in the Mirab Abaya district, Gamo Zone, Southern Ethiopia. The FFL covers an area of about 100 km^2^ and lies between 06°10’ 12” to 06°15’ 00” N latitude and 37°42’ 36” to 37°47’ 24” E longitude with an elevation ranging between 1184-1795 masl (Fig 1).

**Figure 1.**
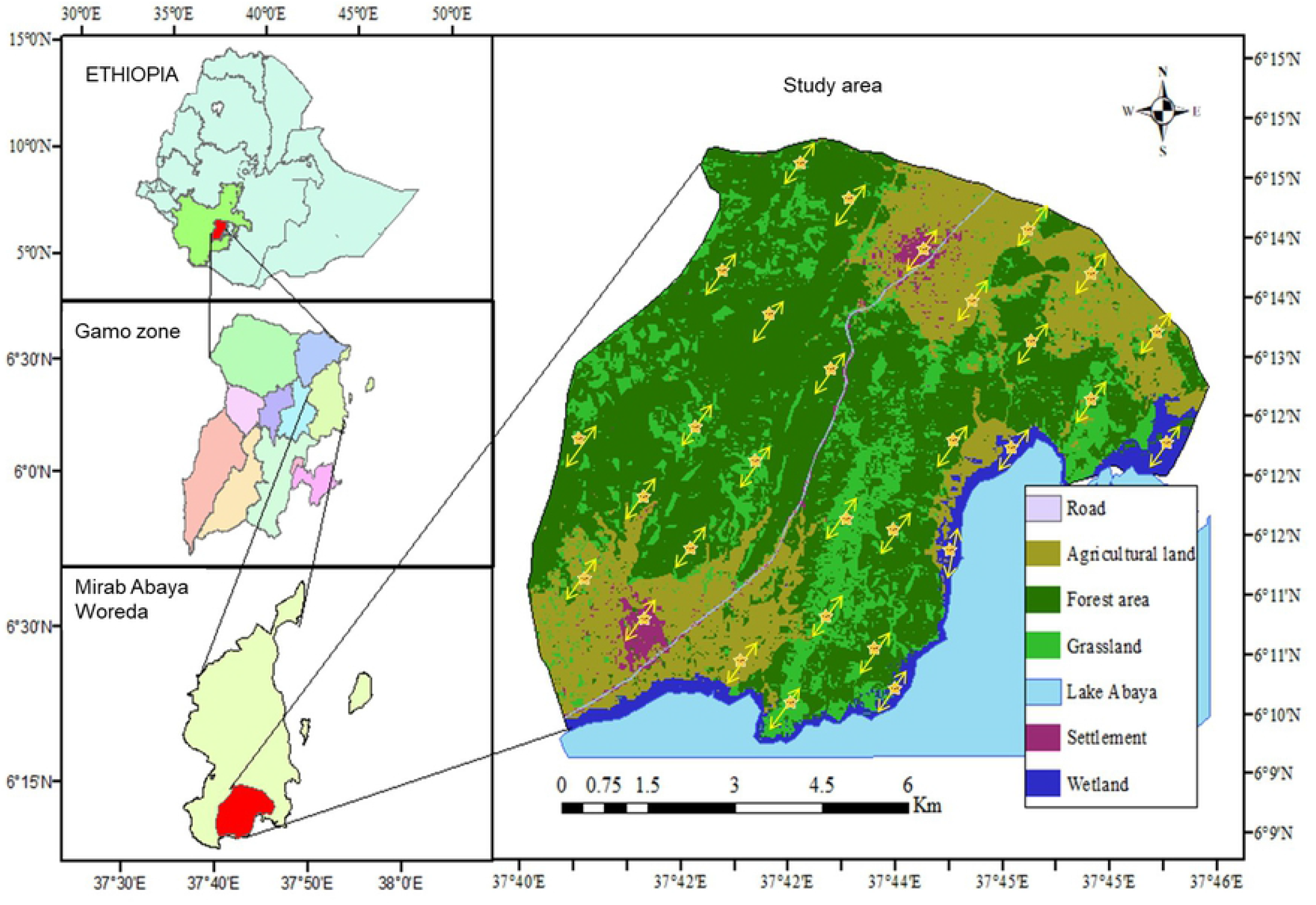
Map of Faragosa-Fura landscape showing land-use types. The arrows indicate transect lines and stars indicate camera points.

The mean monthly temperature and rainfall of the FFL range between 14.75 °C −26.75 °C and 41.8 mm −161.4 mm, respectively [28]. It is bounded by Ankober and Faragosa kebeles (the lowest administrative unit) to the north, Done kebele to the east, and Lake Abaya to the west and southwest, Umo-Lante and Fura kebeles to the south. There is one main asphalt road from Addis Ababa to Arba Minch that crosses the study landscape, making it easily accessible. The landscape is endowed with a wide variety of wildlife species coexisting with humans. Besides, the area is predominantly rural, characterized by settlements and surrounded by agricultural land and forested landscape, and associated with Lake Abaya, the largest lake in the Ethiopian Rift Valley system. Thus, the area is important for wildlife conservation in Ethiopia.

The total population of the area is estimated to be 17,740, of which, 8909 were men and 8831 were women [29]. The main economic activities are farming, livestock breeding, beekeeping, and the collection of forest products. Females are responsible for indoor activities such as caring for children and cooking while males for outdoor activities such as farming and livestock rearing. As a result, the landscape is under the pressure of anthropogenic activities such as agricultural expansion for vegetables and banana plantations, and livestock rearing [30]. The most commonly kept livestock species are cattle, goats, sheep, and chickens.

The common flora observed in the area were Acacia spp, *Terminalia brownie, Dodonaea angustifolia, Acalypha fruticosa, Maytenus arbutifolia, Olea europaea, Ximenia americana, Syzygium guineense, Bridelia scleroneura, Maytenus undata, Vangueria apiculata, Rhus vulgaris*, and *Ozoroa insigns* [30].

### Land-use and landscape drivers

We used a total of seven factors to examine the variation in species richness and distribution of the carnivores in the landscape: six land-use types (forest, wetland, grassland, agricultural land, settlement) and two landscape factors (altitude and distance from the road where pieces of evidence of carnivores recorded). In this study, the heterogeneous landscape was divided into five more homogeneous land-use types during reconnaissance using ArcGIS (Aeronautical Reconnaissance Coverage Geographic Information System). The land-use types designed were forest (area = 36.04 km^2^), wetland (area = 10.12 km^2^), grassland (area = 17.74 km^2^), agricultural land (area = 24.19 km^2^), and settlement (area = 3.23 km^2^). Depending on the size of the survey area, each land-use type is further divided into spatially isolated sites, giving the 30 sites: wetland (4), forest (12), grassland (6), agricultural land (6), and settlement (2) where line transects and camera traps were distributed (Fig 1). In addition, the landscape factors such as altitude and the distance from vehicle road where each piece of evidence was recorded were measured using ArcGIS data layer and Garmin GPS.

### Line-transect design

Line-transect is a well-recognized and cost-effective methodology for surveying different sized vertebrates in tropical forests and savannas [31]. We established, a total of 30 fixed-length transects using each spatially isolated site described above to collect data (Fig 1). The distance between adjacent transects and from habitat edge to a transect was limited to a minimum of 0.5 km, to avoid double counting and to avoid edge effects [31]. The length of each transect was two km and a fixed sighting distance of 100 m on both sides of transects was used in each land-use type.

### Carnivore survey

To gather data along 30 transects and to maximize the survey effort, we employed three complimentary field surveys techniques: Sign survey, camera trapping survey, and opportunistic carnivore sighting survey.

#### Sign survey

fresh tracks, feces, hair, burrows, odor, and digging observed along transects were recorded [32]. Sign survey can enhance the survey efficiency for many mammal species, contributing to maximize the distribution of species lists [33]. To avoid recounting of the same sign during subsequent monthly sampling periods, only the counted signs by data collectors and the researcher was marked at a place.

#### Camera trapping survey

we used infrared digital camera traps (Bushnell Trophy model Cam HD-119447, given a unique ID number, equipped with 32GB SDHC memory cards and eight super alkaline AA batteries, casing, and mounted to a tree with a cable lock), triggered automatically by the animal movement. We distributed cameras equally along transects (see Fig 1 above). We activated cameras with default 10 seconds photographic delay between pictures. We adjusted time, date, month, year, and auto mode for all camera traps. Then, we fixed one camera trap in each transect by prioritizing signs of carnivores. Because the literature revealed that fixing cameras along signs helps to maximize captures within the study sites [16]. We maintained two to three-meter fixing distances on either side from the center of the trail or road to get identifiable photographs and to protect cameras from animal damage [34]. We fixed each camera on the flat ground to maintain a suitable degree, at a height of 30 to 40 cm on the tree above the ground. We faced cameras to the north or south to minimize false triggers from the rising or setting sun, and adjusted parallel to the ground to ensure direct a field of view. Finally, small vegetation such as long grasses and bushes makes it the camera difficult to identify the carnivores cleared and its proper installation checked before leaving the fixed site.

#### Opportunistic sighting survey

all opportunistically observed carnivores counting along transects were done by the naked eye and using Bushnell laser rangefinder binoculars. Opportunistically sighted carnivores killed by vehicles along the asphalted road were also recorded for land-use types.

We collected data from August to September 2020 during the wet season and January and February 2021 during the dry season. We carried out carnivore surveys for three consecutive days per month for four months. We fixed each camera for three days while conducted the transect walk early in the morning between 6:00 and 10:00 hrs.; when most animals become more active) [4, 30]. Therefore, we surveyed each transect line 12 times during the study period. During transect visits, a researcher and trained data collectors walked quietly and gently, and at a constant speed along each transect against the direction of the wind to minimize disturbances of carnivore species. We shifted data collectors between transects to minimize “observer effect (bias)”. We recorded data whenever an individual animal or signs of animals sighted as follows: date, time, land-use type, transect number, species name, body size, altitude, and distance from vehicle road and GPS location.

For photographs and sightings evidence, we used body size, coloration, and dominant behavior to identify carnivore species following the Kingdon Field Guide to African Carnivores [32] and researchers’ field experience. When scat encounters, the field identification of carnivore species was determined based on its morphology, including diameter at the widest point, length, shape, color, odor, and disjoint segments following Chame (2003). Additional criteria include the nature of the scat deposit site, the presence of tracks, den sites, or signs of activity of the species under study [35]. Ambiguous signs were left unrecorded. Data were pooled together for each transect, land-use type, and used for analysis [4]. In addition, the landscape variables such as altitude and distance from vehicle road to each evidence recorded were measured. We oriented data collectors about COVID 19 preventive measures made them maintain social distancing and use face masks throughout the data collection period.

### Data analysis

We used four different statistical tests to identify the differences in species richness and key factors that determine carnivore species richness. These are 1) descriptive statistics for characterizing the mean difference in species richness between land-use types; 2) chi-squared (χ^2^) test for comparing the significance difference in carnivores distribution between 30 sampled sites; 3) ANOVA for assessing the effect of land-use types on both species distribution and richness of carnivores; 4) binary logistic regression for characterizing the effect of landscape factors on both species richness and distribution of carnivores. The value of standardized regression coefficients (β) was used to compare the effect of land-use and landscape factors on species richness (-β = negative relation, + β = positive relation, 0 = no effect). The significant effect of all factors on species richness was examined from each P-value (i.e., the P < 0.05 indicates significance effect) and confidence interval (i.e., CI excludes zero). All statistical significance tests were conducted at an alpha level of 0.05. All statistical analysis was carried out using R 4.0.4 of R development Core Team 2020 (R Core Team, 2020).

## Results

### Species taxonomic and body size composition

Six families composed of 12 species were identified in the FFL (Table 1; SI Fig 1). Family Felidae and Herpestidae were represented by three species each, whereas Family Canidae and Viverridae by two species only. The rest two families were represented by a single species. Out of species identified, two species (*Panthera pardus* and *Crocuta crocuta)* were large-sized, six were medium-sized and the rest were small-sized carnivores (Table 1).

**Table 1.**
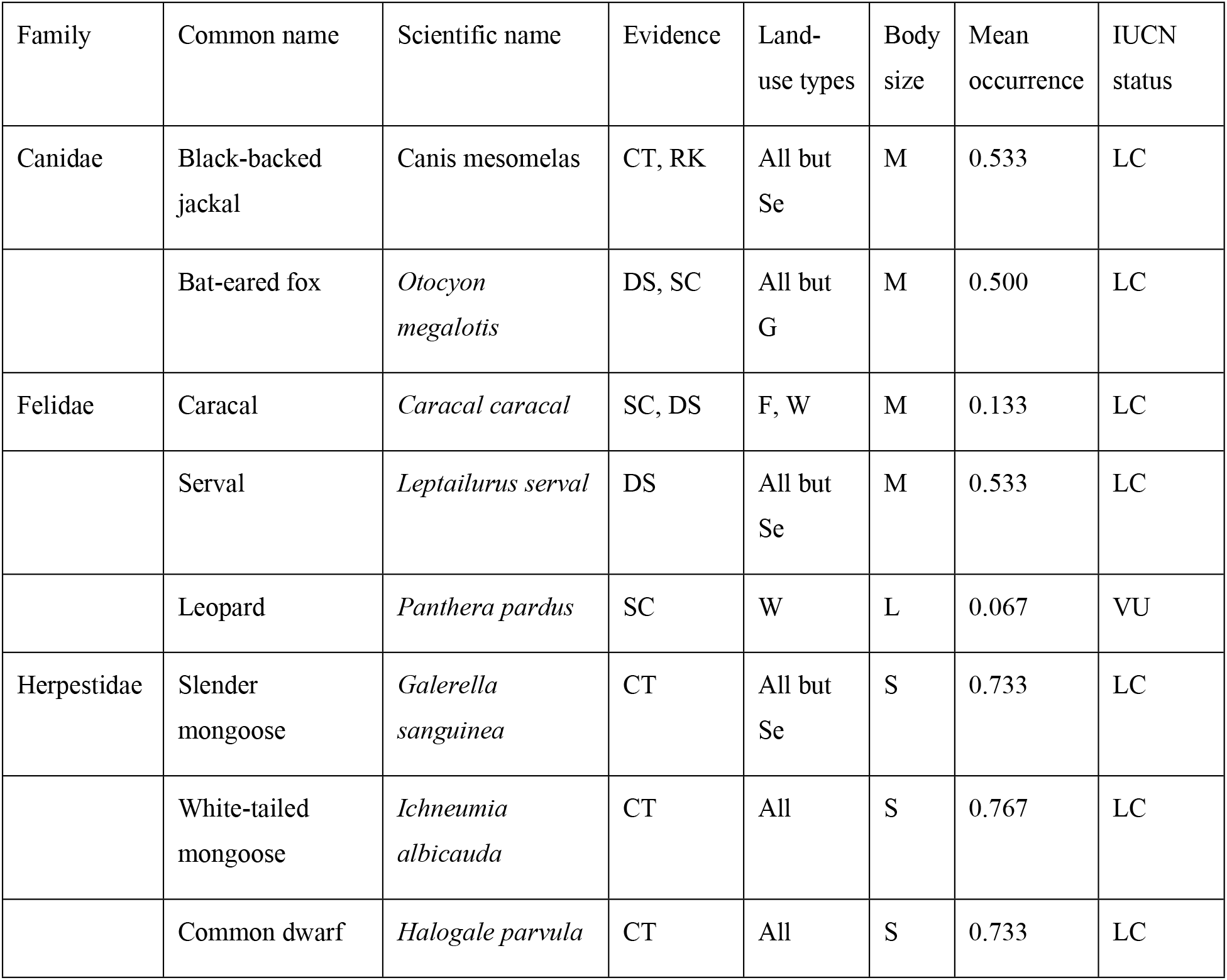
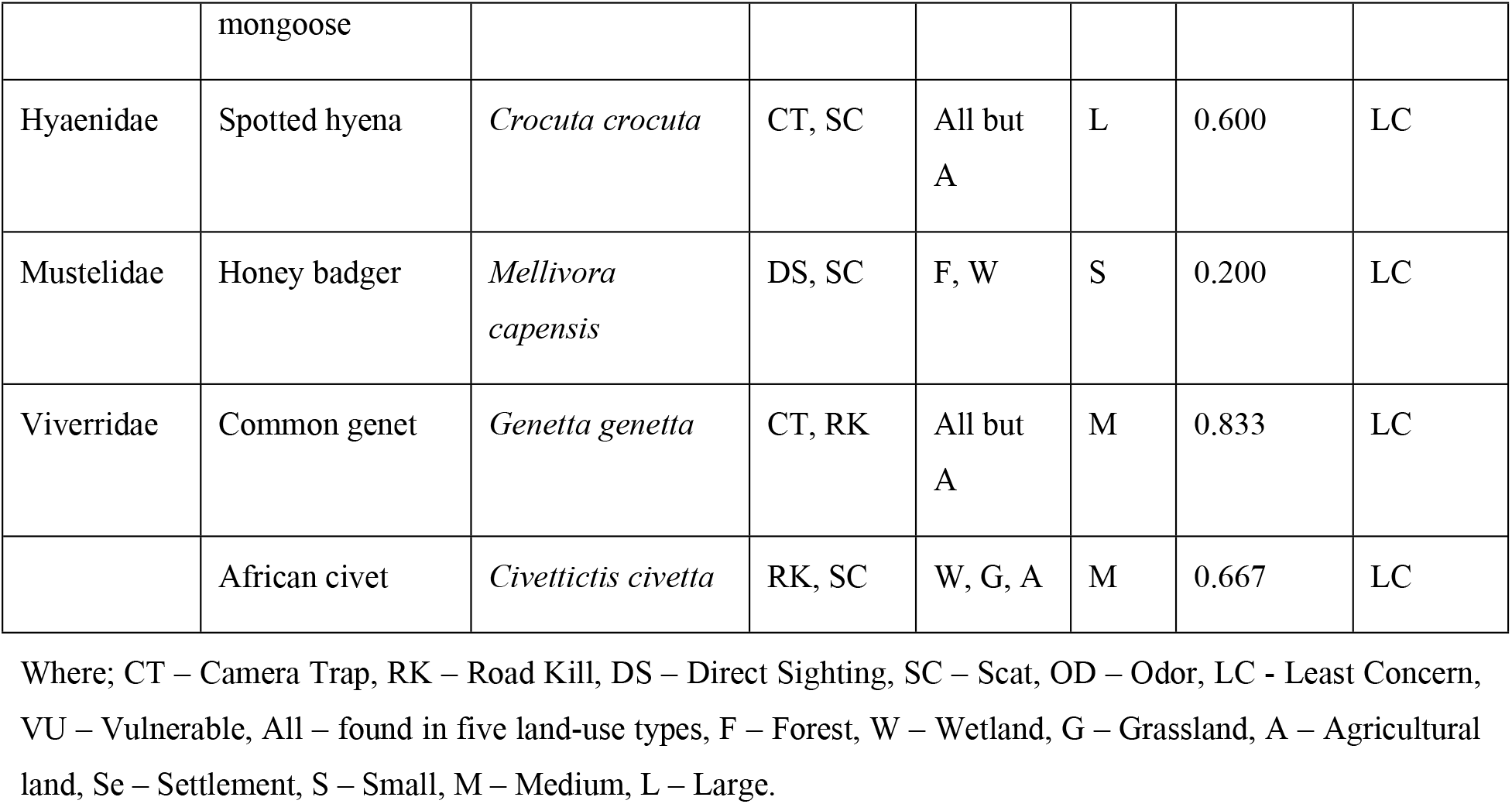
The list of six families and 12 carnivore species identified in the camera trap survey. Their respective IUCN red list category, different names are indicated.

### Species richness and distribution between land-use types

Overall, 12 carnivore species were detected, including the vulnerable Felidae <u>species</u> - *Panthera pardus*. Species richness was highest in the wetland (12) and lowest in settlement (5) (Fig 2). Species richness was varied among the five land-use types types, in increasing order of: settlement (5) < agricultural land (7) < grassland (8) < forest (10) < wetland (12). There was significant difference in species richness between land-use types (χ^2^ = 19.467, df = 4, p = 0.003). The mean species richness was higher in the wetland 7.87±0.494(SE) followed by grassland 6.00±0.548(SE) (Fig 2; SI Table 2). Overall, the mean richness of the study area was 5.73±0.284(SE).

**Figure 2.**
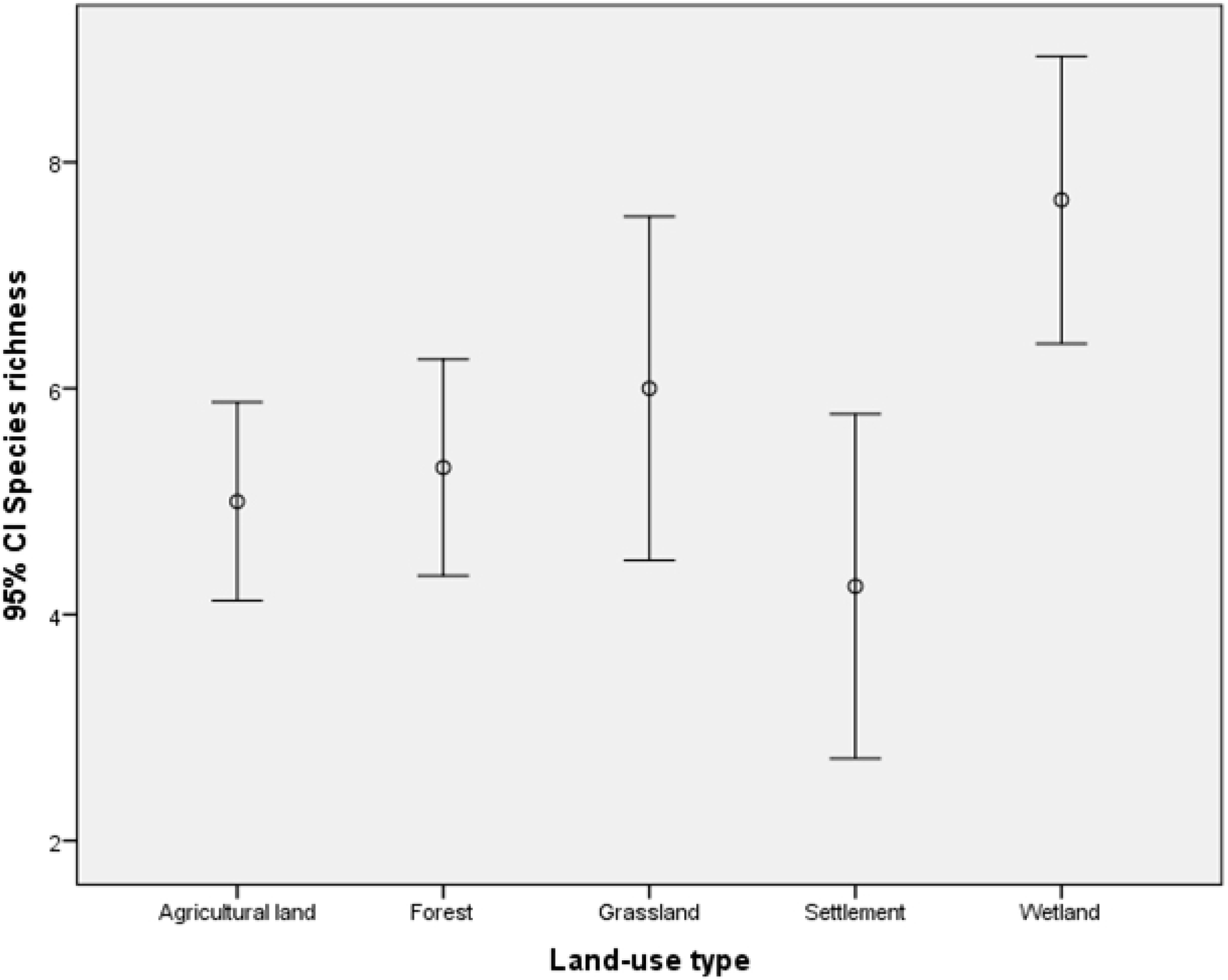
Error bars showing mean carnivore species richness and 95% confidence interval between land-use types.

*Ichneumia albicauda and Halogale parvula* were identified in all land-use types (habitat generalists), while another six species were recorded from four land-use types in common (Table 1). *Panthera pardus* was habitat specialists recorded only in wetland (Table 1). *Mellivora capensis* and *Caracal caracal* were recorded only in the wetland and forest.

Out of 30 sampled sites, *Genetta genetta* was distributed at the 25 sites (the greatest proportion of occurrence = 0.833) followed by *Ichneumia albicauda* (Fig 2). *Mellivora capensis, Caracal caracal*, and *Panthera pardus* were distributed at the fewest sampled sites (Fig 2). *Panthera pardus* was detected only at two sites (the least proportion of occurrence = 0.067).

### Effects of drivers on species richness

Overall, the species richness displayed a negative association with the settlement (Fig 3D) and agricultural land (Fig 3E) whereas exhibited a positive association with other drivers (Fig 2). The species richness exhibited an overall positive and significant association with wetland (β = 0.26; P < 0.01; Fig 3B) and altitude (they prefers higher altitude, β = 0.058, P < 0.05; Fig 3F) and to a lesser extent to distance to road (β = 0.059; P > 0.05, Fig 3G). The weak relationship between drivers and species richness implies that there is variability of occurrence of different species in sampling sites. For example, seven species were positively associated with distance from the road while five species were associated negatively (the overall species richness higher farther from roads, Fig 3G).

**Figure 3.**
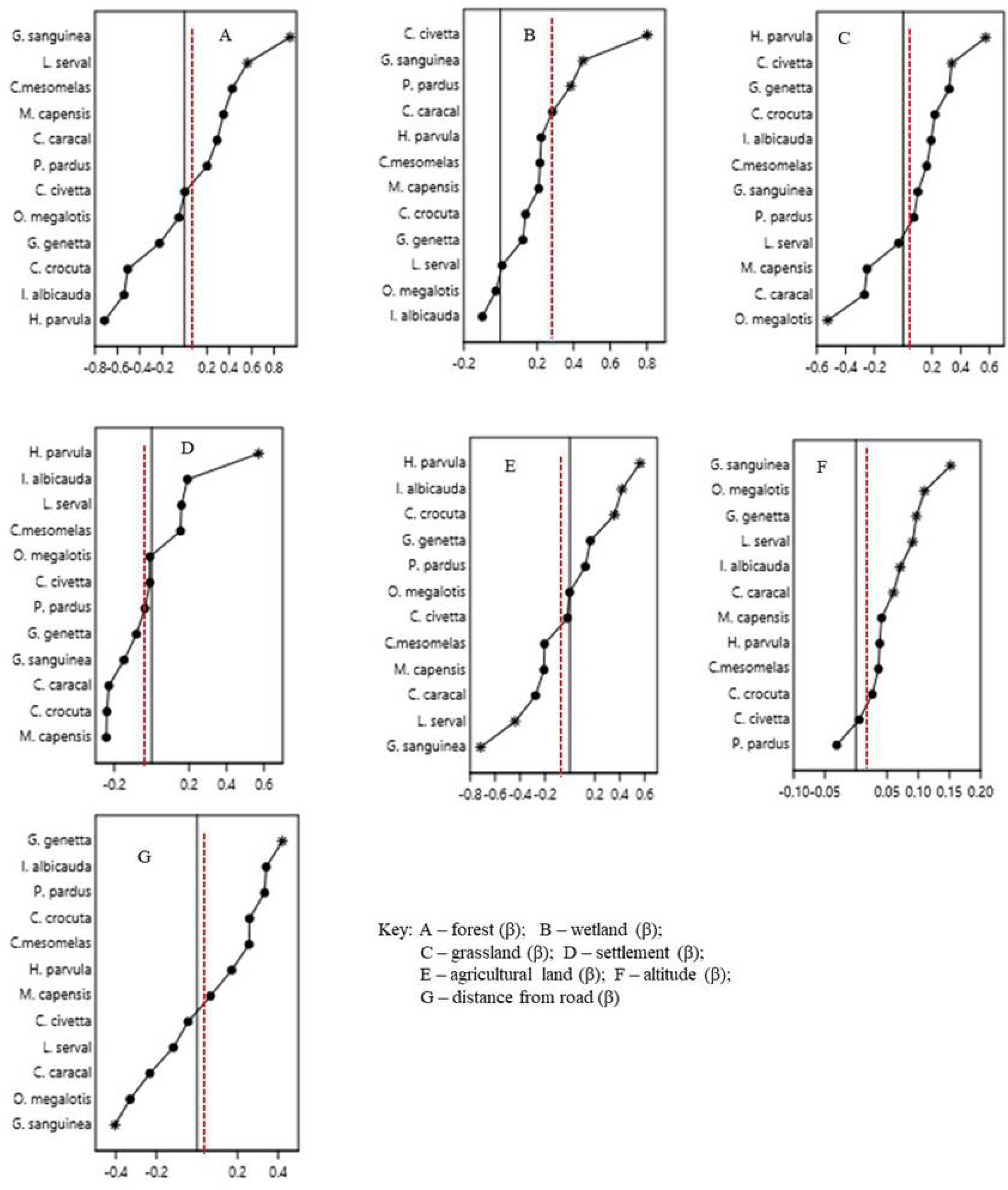
The line plus dot diagram showing the species-specific effect of different drivers on the distribution of carnivore species in the FFL. Stars indicate significant effect (confidence interval exclude zero), dots indicate coefficient (β) values, solid central axis indicates demarcation (zero point) for positive (avoidance) and negative association (preference), red dot line indicate community effect.

### Effects of drivers on the specific-species

Seven out of 12 species displayed positive association with forest, and this association was significant for two species: *Galerella sanguinea* (β = 0.562; P < 0.001) and *Leptailurus serval* (β = 0.943; P < 0.05; Fig 3A; SI Table 1). Among 10 species showed positive association with wetland, *Civettictis civetta* (β = 0.803; P < 0.001), *Galerella sanguinea* (β = 0.459; P = 0.01) and *Panthera pardus* (β = 0.384; P < 0.05) displayed the significant association (Fig 3B; SI Table 1). Out of eight species displayed positive association with grassland (Fig 3C; SI Table 1). The association was significant for *Halogale parvula* (β = 0.573; P < 0.01) and *Civettictis civetta* (β = 0.337; P < 0.001) whereas four species displayed negative association with grassland and the association was significant only for *Otocyon megalotis* (β = −0.524; P < 0.001).

Four and eight species associated with settlement positively and negatively, respectively (Fig 3D; SI Table 3). Only *Halogale parvula* (β = 0.572; P < 0.01) was strongly and positively associated with settlement. Five species showed positive association and seven species showed a negative association with agricultural land (Fig 3E; SI Table 3). *Halogale parvula* (β = 0.567; P < 0.01), *Ichneumia albicauda* (β = 0.417; P < 0.05) and *Crocuta crocuta* (β = 0.357; P < 0.05) displayed significant positive association whereas *Leptailurus serval* (β = −0.439; P < 0.05) and *Galerella sanguinea* (β = −0.716; P < 0.001) displayed significant negative association with agricultural land.

Six species displayed positive association with altitude, of which five showed a significant preference to the higher altitude (Fig 3F; SI Table 3). The five species that showed strong preference to higher altitude were *Galerella sanguinea* (β = 0.152; P < 0.001), *Otocyon megalotis* (β = 0.110; P < 0.001), *Genetta genetta* (β = 0.097; P < 0.01), *Leptailurus serval* (β = 0.091; P < 0.01), *Ichneumia albicauda* (β = 0.071; P < 0.05) and *Caracal caracal* (β = 0.060; P < 0.05). Although six species showed a negative association with altitude, none of them exhibited significant association (P > 0.05; Fig 3F; SI Table 3)). The carnivore species association with distance from the road was variable: seven out of 12 species showed positive association (prefers farther from the road) while five species showed a negative association (Fig 3G; SI Table 3). *Genetta genetta* (β = 0.405; P < 0.001) and *Galerella sanguinea* (β = −0.421; P < 0.001) significantly positively and negatively associated with road, respectively. *Canis mesomelas and Mellivora capensis* were the two species not significantly affected by any of drivers studied (Fig 2; SI Table 3).

## Discussion

Using multiple survey techniques enhances survey efforts and provides insight into the first quantitative data at the community level of carnivores. Most surveys in Africa focus on single species [16]. In addition, the result indicates that the coexistence of humans and carnivores is possible in a human-dominated landscape in the study area. Several earlier studies have also shown humans and carnivores coexist [6, 37, 38]. Since little is known about carnivore distribution in Ethiopia or more generally across much of East Africa [2, 39], this survey confirms that FFL harbors at least 12 carnivore species, including the global conservation concern *Panthera pardus*, and is one of the most important areas for the conservation of carnivores in Ethiopia. The coexistence with carnivores appears to foster the development of management tools.

### Taxonomic composition and species richness

All the six families of carnivores recognized earlier in Ethiopia (Lavrenchenko & Bekele, 2017) are also recorded by this study. Durant et al. (2010) identified six families of carnivores in the Serengeti ecosystem, Tanzania [15]. In the present study, Felidae and Herpestidae were composed of three species, while Hyaenidae and Mustelidae were each composed of a single species. The richness of Herpestidae (mongooses) could be due to their more adaptive nature to different land-use types, diversified foraging behavior (fruits, meat), and high tolerance level to human disturbances [7, 20, 39]. For the Felidae, it could be wild and domestic prey availability, high vegetation cover, and access to water [14, 34]. The low species richness of the family Hyaenidae and Mustelidae is might be associated with anthropogenic pressure in the area.

Out of 32 carnivore species of Ethiopia [2, 3], this study documented 12 carnivore species. Literature proved that carnivores are crucial in regulating and maintaining habitats [11]. Out of 12 carnivore species, two were large-sized – *Crocuta crocuta* and the global conservation concern species vulnerable *Panthera pardus* [5]. Large carnivores in particular are often used as “keystone species” in conservation efforts, thereby creating more resilient habitats [12]. Also, studies proved that areas containing conservation concern species are important for conservation practice [8, 11, 33]. Thus, the FFL is one of the most important areas for the diversity of carnivores in Ethiopia. This has important implications for carnivore conservation and management.

These multiple surveys provided the highest number of carnivore species compared to indirect sign survey and direct visual survey in different localities in Ethiopia. Qufa and Bekele identified four species of carnivores from the Lebu Natural Protected Forest, Southwest Shewa, Ethiopia [33]; and Girma and Worku identified six species of carnivores in the Nensebo Forest, Southern Ethiopia [4], which is lower than the present study. The higher species composition and richness of carnivores in the present study area might be associated with the inclusion of advanced survey technology (camera trapping), a higher survey period, access to water and vegetation cover.

### Effect of land-use types on carnivores

As expected, wetland had the highest species richness and almost all (83.3%) carnivore species were positively associated with wetland (significant for *Civetta civetta, Galerella sanguinea*, and *Panthera pardus*) (Fig 2). Given the smaller area (only 10.12 km^2^) of the wetland compared with the forest area (36.04 km^2^), the highest species richness and strong positive preference of most species are surprising and against the “area-species relationship”. Habitats with greater area tend to contain a higher number of species compared with habitats with a smaller area [10]. In addition, the wetland supports unique species, specifically the vulnerable *Panthera pardus*. Possibly this is because of where preys were easier to catch during watering. The wetland provides a wide array of provisioning, supporting, cultural, and regulating services contributing to wildlife [18]. Prior studies have noted the importance of water and cover as a major habitat requirement [11]. Thus, the presence of conservation concern species, strong preference of most species, and the highest species richness in the wetland demonstrate that the wetland is playing an important role in wildlife conservation in the FFL. This might be due to access to water (Lake Abaya), dense forest, and less anthropogenic pressure.

Forest is the second most abundant in terms of species richness (contains 10 species) and above two-thirds of carnivores were positively associated with it. This is possibly due to high vegetation cover, cooling effect, and exerts less human disturbances. Observed associations between carnivore species agree with previous research in Ethiopia on *Panthera leo* [41], *Crocuta crocuta* [42], and *Civettictis civetta* [24]. This highlights the importance of forest in maintaining functional assemblage and/ or diversity, and ecological integrity besides provision of the basic need such as food, shelter, and cover [18]. Worldwide, studies reported that the forest is the most important habitat for wildlife and biodiversity conservation [3, 5, 11].

Although grassland is open and unable to hide elusive carnivore species, it contains eight species and a large number of carnivore species were positively associated with it (significant for *Halogale parvula, Civettictis civetta, Genetta genetta*). Possibly they might have been attracted to grassland due to promote ease of movement, access to termites and insects, anti-predator scanning, and prey availability [4, 18]. Therefore, the land-use types should be given equivalent conservation attention. Further, focused studies are needed on prey-predator relationships for effective management planning in the FFL.

Studies proved that species richness is likely to be lower in the settlement and agricultural lands due to increased anthropogenic activities such as farming, poaching, and controlling carnivores by killing not to damage livestock [7, 39]. As expected, this is supported by our finding that the settlement (five species) and agricultural land (seven species) had fewer carnivore species than other land-use types. In addition, Congruent with the hypothesis of the present study, most of the carnivores species would negatively associate with agricultural land. However, this generalization varies between species and probably includes aspects of dispersal, colonization, and the tolerance of stress [39, 43, 44]. For example, the positive associations were significant only for *Halogale parvula, Ichneumia albicauda*, and *Crocuta crocuta*, this might be due to access to domestic prey for three of them, scavenging (*Crocuta crocuta*) and omnivores behaviors (*Halogale parvula, Ichneumia albicauda*). The present result, perhaps indicative of an “ecological trap” (i.e. environmental change leads organisms to prefer to settle in poor-quality habitats; Ripple et al., 2014). However, these species are known for livestock predation and crop damage [45], and thus, agriculture development could lead to decreased habitat and isolated population [43] and may escalate the conflict between human and carnivores, which is a threat to biodiversity conservation. Thus, land sharing (wildlife-friendly agricultural system) for agricultural adapted carnivore species should be implemented to consider the conservation of carnivores[46].

Although the effect was statistically weak, eight out of 12 carnivore species had displayed a negative association with the human settlement [39], which might be due to hunting in retaliation for livestock depredation and crop damage [15, 17, 45]. These patterns are shared across East Africa and worldwide, where carnivore avoids human disturbance except a few members (e.g., hyena) that are known to attack livestock inside human settlements at night and attracted by anthropogenic food sources (e.g., mongooses) [20, 39]. Similarly, previous studies suggest human disturbance negatively affects carnivores in southern Ethiopia [18, 41, 44]. The only *Halogale parvula* was a positive and significant association with the settlement, probably due to the high density of rodents and fruit trees that may provide easy pickings for this omnivorous species.

### Effects of landscape factors on carnivores

Carnivore species richness increased significantly with altitude. Overall, the higher altitude had a significant effect on half of the carnivore species (*Galerella sanguinea, Otocyon megalotis, Genetta genetta, Leptailurus serval, Ichneumia albicauda, Caracal caracal)*. Except for the *Panthera pardus*, all carnivore species were positively associated with altitude. This is likely due to higher altitude sites are far away from human habitation (thus exerts less disturbance), prey cascading (scanning), and low temperature at higher altitudes. Higher temperatures result in habitat shifts mostly toward upper elevation [15]. This agrees with studies by other scholars elsewhere [39].

The species richness was higher, farther from vehicle road; this might be from road-kill, collisions with vehicles, and human disturbance. Although only one species (*Galerella sanguinea*) displayed a strong negative response, the majority tend to avoid roads (Fig 2). The effect of roads on carnivores has been reported elsewhere [1]. The vehicle kills to genet, African civet, and jackal were recorded during the study period. Penetrating natural forests with roads results in opening avenues for human encroachment, road-kill, and species collisions with vehicles [1,19]. Roads, in general, have been associated with cascading effects like overexploitation, habitat conversion, fire, farming, and invasive species [1,19].

### Conclusion and conservation implications

This study provides information on the differences in species richness and distribution pattern of carnivores across land-use types of hitherto little known human-dominated landscape. The findings of the study revealed that local people coexisting with 12 carnivore species, including globally vulnerable *Panthera pardus* [5]. The study also attempted to collect ecological information on the species richness and distribution of carnivores using multiple surveys (sign, camera-trapping technology, direct sighting), which would serve as valuable baseline information for stakeholders to make imperative conservation decisions and for researchers wishing to conduct related ecological studies in a human-dominated landscape.

The wetland is the most important land-use type of species because of water availability and less human pressure; provides a tangible target for carnivore conservation in the FFL. Forest affected species richness, it is possible that carnivore species persisting in the forest have proven themselves more resilient to human impacts like hunting, having passed through the “extinction filter” that claimed other species [12]. A considerable amount of species prefers grassland, which might be due to promoting ease of movement, access to termites, and insects, anti-predator scanning, and prey availability. Anthropogenic factors such as agriculture, settlement, and vehicle roads are influential factors affecting carnivore species significantly.

As expected, many of the carnivore species sampled by the present study require largely less disturbed and forested habitat and may be seriously ill-adapted to the agricultural land, settlement, and vehicle roads. Therefore, intensifying agricultural production on lands already converted, while prioritizing the protection from the conversion of native habitats is in need. Considering the likelihood that wild habitat will decline due to agricultural activity, in Ethiopia, as elsewhere, both the protection of ecologically rich areas (land-sparing-segregating wildlife conservation from agricultural production) and land-sharing (wildlife-friendly agricultural system) to conserve wildlife are crucially important [9, 46]. The federal and regional governments should legalize the FFL as a wildlife refuge area to conserve the species of the area.

## Acknowledgments

Our special gratitude goes to the Mirab Abaya Wereda Administrative office for allowing us to research in the Faragosa-Fura landscape. We also duly acknowledge Faragosa and Fura Kebeles’ administrative office and agricultural extension workers for their assistance during data collection in the habitat types. We also thank the Department of Biology, College of Natural Science, Arba Minch University, for their invaluable logistics and financial support. We are thankful to the Arba Minch College of Teacher education for the logistic support as well as covering the living and other costs of the Ph.D. candidate and we are grateful for the support provided.

## Availability of data

All data used are included in the article and supplementary material.

## Author Contributions

All authors conceived and designed the data collection procedures. The first author conducted the household survey, analyzed, wrote up the manuscript, and revised the whole document. The second and third authors designed the survey method, supervised the survey process, edited the manuscript, and revised the final version of the main document for submission for potential review. All authors approved the submitted version.

## Permits

The journal shall own the work such as copyright, the right to republish the article in whole or in part for sale or free distribution, the right to translate them into languages other than English, and the right to republish the work in a collection of articles in any other mechanical or electronic format.

